# A persistent intracellular bridge and cell cycle misregulation enable polar body cell divisions and tumor formation in *Mos*-deficient eggs

**DOI:** 10.1101/2025.08.26.672293

**Authors:** Gisela Cairo, Muhammad A. Haseeb, Karen Schindler, Soni Lacefield

## Abstract

Mammalian female meiosis is uniquely regulated to produce a developmentally competent egg capable of supporting embryogenesis. During meiosis I, homologous chromosomes segregate, with half extruded into the first polar body. The egg then arrests at metaphase II and only resumes meiosis and extrudes the second polar body following fertilization. The MOS/MAPK signaling pathway is important for maintaining the metaphase II arrest; in *mos^-/-^* mutants, a subset of eggs undergo spontaneous parthenogenetic activation and exhibit additional abnormal cell divisions. To further understand the cell cycle mis-regulation in activated *mos^-/-^* eggs, we used time-lapse microscopy to monitor the abnormal divisions. We discovered that, following parthenogenetic activation, the first polar body can assemble a spindle, segregate chromosomes, and divide with timings similar to anaphase II onset in the egg. This behavior contrasts with wildtype polar bodies, which do not divide and are typically degenerated. We demonstrate that *mos^-/-^* eggs and polar bodies can exchange cytoplasm at the time of meiosis II spindle assembly, likely allowing the transfer of cell cycle regulators between the two compartments. Further inspection revealed that *mos^-/-^* eggs have defective meiotic midbody assembly with most eggs lacking a cap structure, which is needed to separate the two compartments. We report that polar bodies of *mos^-/-^* eggs can re-enter the cell cycle and undergo additional aberrant divisions. These findings identify MOS as a critical regulator of meiotic midbody formation and uncover a novel consequence of disrupted MOS/MAPK signaling: the potential for polar bodies to become mitotically active and contribute to tumor formation.

## INTRODUCTION

Meiotic chromosome segregation is tightly regulated during mammalian oogenesis to ultimately produce an egg capable of supporting early embryonic development. Meiosis is initiated in the fetal ovary with DNA replication, programmed double strand break formation and repair, followed by a long prophase I arrest. At sexual maturation, a subset of oocytes resume meiosis and enter the first meiotic division. A bipolar spindle assembles and localizes to the cortex, resulting in an asymmetric division in which half the homologous chromosomes are retained in the oocyte and half are extruded into a smaller polar body. In telophase I, the actomyosin ring contracts and a specialized structure known as the meiotic midbody assembles, recruiting proteins needed for abscission during cytokinesis (Jung et al., 2023; Uraji et al., 2018). The oocyte then assembles a bipolar spindle and arrests at metaphase II, awaiting fertilization (Mihajlović and FitzHarris, 2018). The polar body eventually degenerates.

The meiotic divisions are orchestrated by the dynamic interplay between cyclin-dependent kinase 1 (CDK1) and the Anaphase Promoting Complex/Cyclosome (APC/C), an E3 ubiquitin ligase (Kim et al., 2023; Pines, 2011). CDK1, in complex with a B-type cyclin, phosphorylates substrates essential for meiotic progression, including those involved in spindle assembly and chromosome alignment at the metaphase plate. CDK1 activity also promotes activation of the APC/C, which in turn ubiquitinates cyclin B and other regulatory proteins, targeting them for subsequent proteasomal degradation, triggering anaphase I onset. To enter meiosis II, APC/C activity must be inhibited, allowing cyclin B1 levels to reaccumulate and sustain high CDK1 activity. The metaphase II arrest is maintained by the sustained CDK1 activity in conjunction with inhibition of APC/C (Hörmanseder et al., 2013).

In female mammals, establishment of APC/C inhibition requires EMI2 (Early Meiotic Inhibitor 2), while maintenance of the metaphase II arrest requires both EMI2 and MOS (Moloney Sarcoma Oncogene) (Hörmanseder et al., 2013). EMI2 functions as a direct inhibitor of APC/C, whereas MOS, a serine/threonine kinase, is thought to maintain EMI2 stability and indirect inhibition of APC/C through activation of downstream kinases. Although originally identified as the transforming gene of Moloney murine sarcoma virus (*v-mos*), *Mos* expression is largely restricted to oocytes (Dupré et al., 2011). MOS is the upstream activator of the MAPK cascade (mitogen-activated protein kinase), which is essential for sustaining metaphase II arrest. Notably, a subset of eggs from *mos^-/-^* mice fail to maintain this arrest, and instead undergo spontaneous parthenogenetic activation and premature anaphase II (Cairo et al., 2025; Colledge et al., 1994; Hashimoto et al., 1994; Verlhac et al., 2000a). Some of these activated eggs proceed through additional abnormal cell divisions. Approximately one third of female *mos^-/-^*mice develop ovarian germ cell tumors, suggesting that the activated eggs that escape ovulation can proliferate aberrantly in the ovary (Colledge et al., 1994; Hashimoto et al., 1994).

We sought to determine the molecular and cellular causes of the abnormal cell divisions observed in *mos^-/-^* eggs. Using time-lapse imaging, we found that in parthenogenetically activated *mos^-/-^* eggs, the first polar body can assemble a spindle, segregate chromosomes, and undergo cell division with timing similar to the inappropriate onset of anaphase II in the egg. These findings were unexpected, as polar bodies in wildtype eggs are typically destined for degeneration. Additionally, we observed that the cytoplasm of the *mos^-/-^* egg and polar body remains connected for an extended period, allowing cytoplasmic components to pass between the two compartments. Further investigation revealed that this prolonged connection is likely due to defective formation of the meiotic midbody, leaving an open intracellular bridge between the egg and the polar body. Notably, polar bodies can re-enter the cell cycle, implicating them as potential contributors to tumor formation. Together, these findings identify MOS as a key regulator of meiotic midbody assembly and demonstrate its role in preventing abnormal divisions in both the egg and polar bodies.

## RESULTS

### Polar bodies undergo chromosome segregation and cell division in *mos^-/-^* eggs

Previous reports showed that *mos^-/-^* mutant eggs undergo a variety of phenotypes: some extrude one polar body, others extrude two polar bodies, and some undergo additional aberrant divisions after anaphase II (Cairo et al., 2025; Colledge et al., 1994; Hashimoto et al., 1994; Verlhac et al., 2000a). In addition, a previous study reported that polar bodies of *mos^-/-^* eggs underwent abnormal cleavage events, unlike wildtype eggs whose polar bodies degenerated (Choi et al., 1996). To gain further insight into these aberrant cleavage events and their potential contribution to abnormal cell divisions, we performed time-lapse imaging of wildtype and *mos^-/-^* eggs. We collected oocytes at prophase I arrest (germinal vesicle; GV), released them from the arrest, and then monitored the resumption of meiosis. We initiated the imaging between 13-15 hours after prophase I release and then imaged for 24 hours so that we could monitor meiosis II and beyond (Figure 1A). We added SiR-DNA and SPY555-tubulin to visualize the chromatin and spindle, respectively. Wildtype cells typically remained arrested at metaphase II and the polar body eventually degenerated (Figure 1B). To our surprise, we found that 53% of *mos^-/-^* eggs formed bipolar spindles in the first polar body (PB1), which elongated and segregated chromosomes (Figure 1C-E). We also observed bipolar spindle formation in *mos^-/-^* eggs in our immunofluorescence images of fixed *mos^-/-^* eggs, suggesting that the phenotypes observed were not due to phototoxicity or the use of dyes (Figure S1A).

**Figure 1.**
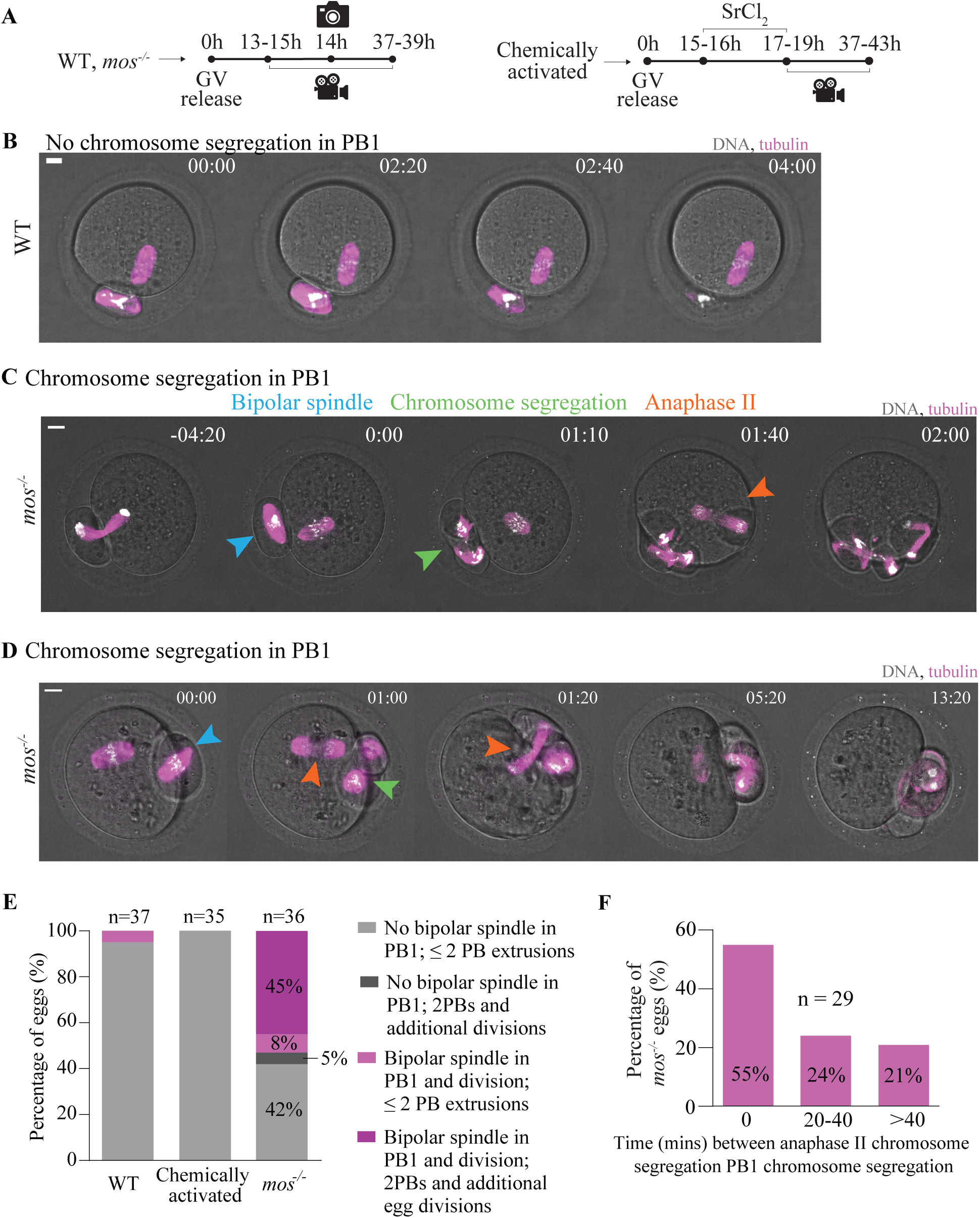
Chromosomes segregate inside polar bodies of *mos^-/-^* eggs with similar timing as anaphase II onset. **(A)** Schematic of experiment outline. **(B)** Representative time-lapse images of wildtype eggs. **(C-D)** Representative time-lapse images of *mos^-/-^* eggs undergoing chromosome segregation events in the first extruded PB/mass. SPY555tubulin and SirDNA was used to visualize tubulin (magenta) and DNA (white), respectively. Time 0 represents the moment of bipolar spindle formation inside the first extruded PB. Blue arrowheads indicate bipolar spindle formation inside the PB; orange arrowheads indicate anaphase II in the egg; green arrowheads indicate chromosome segregation inside the PB. **(E)** Percentage of wildtype, chemically activated, and *mos^-/-^* eggs that exhibit bipolar spindle formation and PB division as well as the number of masses extruded. n indicates the number of eggs imaged per condition. 3 or more independent experiments were conducted per condition. **(F)** Percentage of *mos^-/-^* eggs with polar body division and anaphase II with time intervals of 0 minutes, 20-40 minutes, or more than 40 minutes between events (n indicates the number of eggs imaged within the time interval. 3 or more independent experiments were conducted). **(B-C)** Scale bars, 10µm. GV, germinal vesicle; WT, wildtype; PB1, first extruded polar body; SrCl_2_, Strontium Chloride.

We further categorized the *mos^-/-^* eggs with polar body divisions by their phenotypic outcomes: whether they underwent a second polar body extrusion and additional divisions beyond anaphase II. Most of the *mos^-/-^* eggs with polar body divisions underwent anaphase II, extruded the second polar body (PB2), and then continued to undergo additional divisions (45% of total *mos^-/-^* eggs) (Figure 1E). A smaller fraction of *mos^-/-^* eggs underwent anaphase II to make a second polar body but did not proceed to undergo additional divisions (8% of total *mos^-/-^* eggs). Only 5% of *mos^-/-^* eggs extruded two polar bodies and underwent additional divisions but did not divide the first polar body (Figure 1E). Therefore, most *mos^-/-^*eggs that parthenogenetically activated and underwent additional divisions also had polar bodies that divided.

In support of a previous study, we also found that fewer *mos^-/-^*eggs had polar bodies that degenerated in comparison to wildtype eggs (Figure S1B-D) (Choi et al., 1996). During the time of our imaging, that spanned approximately 24 hours after the oocytes underwent anaphase I, 50% of polar bodies from wildtype eggs and 25% of polar bodies from *mos^-/-^* eggs degenerated (Figure 1B, S1B-D). Because approximately 47% of eggs did not form spindles in the polar body and only 25% of the polar bodies degenerated, these results suggested that some of the polar bodies that did not divide in *mos^-/-^* eggs were also not degenerated or had delayed degeneration (Figure 1E, S1D).

The polar bodies of wildtype eggs do occasionally have abnormal membrane invaginations that are quite different than the polar body divisions observed in *mos^-/-^* eggs. Approximately 20% of polar bodies of wildtype eggs underwent abnormal membrane cleavage, but most did not form bipolar spindles, nor did they segregate chromosomes in the polar body. (Figure S2A-C). Instead, the abnormal membrane cleavage occurred without other noticeable cell cycle events (Figure S2B). All *mos^-/-^* eggs that divided their polar bodies assembled a bipolar spindle and segregated chromosomes in the polar body (Figure S2C, 1B-D).

We next asked if bipolar spindle assembly in the polar body was a common feature of parthenogenetic activation. To address this question, we used strontium chloride (SrCl_2_) to chemically activate wildtype eggs to undergo parthenogenesis (Figure 1A) (Jellerette et al., 2000). The chemical activation triggers processes that occur during fertilization, such as maternal mRNA processing, increased protein synthesis, mitochondrial activity for ATP production, and membrane dynamics such as cortical granule exocytosis (Bellido-Quispe et al., 2024; Horner and Wolfner, 2008; Jellerette et al., 2000; Krauchunas and Wolfner, 2013). Upon chemical activation, eggs undergo anaphase II, extrude a second polar body, and initiate embryo-like divisions. We found that polar bodies in chemically activated eggs did not assemble bipolar spindles; nor did they segregate chromosomes (Figure 1E, S2C). Therefore, the feature of polar body chromosome segregation and cell division is unique to *mos^-/-^*eggs, and not to eggs that undergo parthenogenetic activation.

### Polar body divisions occur with similar timings as anaphase II onset in *mos^-/-^* eggs

In our time-lapse imaging, we noticed that *mos^-/-^* eggs often underwent polar body divisions with similar timings as anaphase II onset and the second polar body extrusion. To quantify this observation, we monitored the time between anaphase II onset and polar body division (Figure 1F). 55% of *mos^-/-^* eggs underwent both events simultaneously (Figure 1D, 1F). In the remaining *mos^-/-^* eggs with polar body divisions, there was a period of either 20-40 minutes (24%) or greater than 40 minutes (21%) between egg and polar body divisions (Figure 1C, 1F, 2A). In some *mos^-/-^*eggs, anaphase II onset occurred prior to the polar body division (Figure 2A) and in others, the polar body division occurred prior to anaphase II onset (Figure 1C).

**Figure 2.**
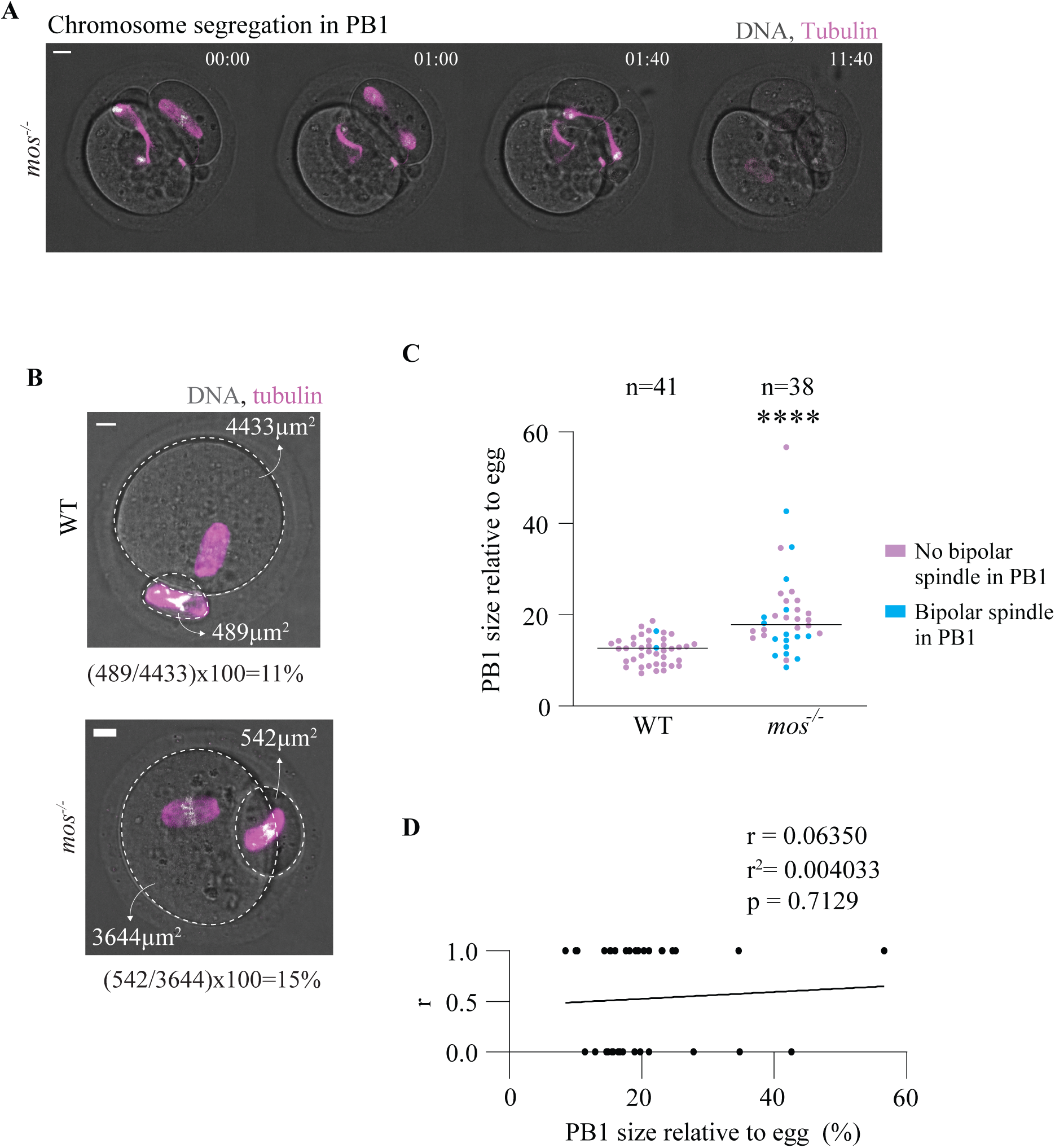
Polar body size does not correlate with its ability to divide in *mos^-/-^* mutants. **(A)** Representative time-lapse images of a *mos^-/-^* egg with a large polar body. Wildtype and *mos^-/-^* oocytes were imaged live 13-15h after prophase I release for 24 hours. **(B)**. Representative time-lapse images of wildtype (top) and *mos^-/-^*eggs (bottom) after the first PB extrusion with mass of the egg and PB and calculation of the PB size relative to the egg. SPY555tubulin and SirDNA were added to visualize tubulin (magenta) and DNA (white), respectively. Scale bars, 10µm. **(C)** Graph showing the size of the first PB extruded relative to the egg in WT and *mos^-/-^* eggs. n indicates the number of eggs imaged per condition. 3 or more independent experiments were conducted per condition. Asterisks indicate a statistically significant difference compared with chemically activated eggs (****, P < 0.0001, Mann-Whitney test). **(D)** Pearson correlation graph. The Y axis represents Pearson correlation coefficient and the X axis represents PB1 size relative to egg. **(A-D)** WT, wildtype; PB1, first extruded polar body; r, correlation coefficient.

We found the result that most *mos^-/-^* eggs divide their polar body with timing similar to anaphase II onset surprising because we did not expect the two compartments to maintain cell cycle coordination. We propose two hypotheses that explain how the two compartments could maintain coordination that we test in the following sections: i) large polar bodies in *mos^-/-^* eggs could contain more cell cycle regulators that allow cell division; or, ii) the egg and the polar body maintain a cytoplasmic connection and share cell cycle regulators.

### Polar body size does not correlate with polar body divisions

Previous studies reported that polar bodies are larger in *mos^-/-^*eggs compared to wildtype. This increase in size is due to a failure to position the spindle close to the cortex prior to division (Choi et al., 1996; Verlhac et al., 1996; Verlhac et al., 2000a). We also observed larger polar bodies in some of the *mos^-/-^* eggs in our time-lapse imaging (Figure 2A). We hypothesized that a larger polar body could allow for more cytoplasm and cell cycle regulators to enter the polar body, allowing a coordinated cell division. We therefore asked if the larger polar bodies were more likely to assemble a spindle and divide than the normal-sized polar bodies. We measured the size of the first extruded polar body relative to the egg by calculating the polar body areas as a percentage of the egg area in wildtype and *mos^-/-^*eggs (Figure 2B). We found that the polar bodies were, on average, larger in *mos^-/-^*eggs compared to wildtype, supporting previous studies (Figure 2C) (Choi et al., 1996; Verlhac et al., 1996; Verlhac et al., 2000a). However, we found that both large and normal-sized polar bodies in *mos^-/-^*eggs assembled bipolar spindles and divided (Figure 2C). Similarly, there were both large and normal-sized polar bodies that did not assemble spindles and divide. Using Pearson correlation, we found no significant correlation between the size of the polar body and whether they divided (correlation coefficient r=0.06; Figure 2D). In conclusion, our findings do not support the hypothesis that larger polar bodies are more likely to undergo division than normal-sized polar bodies.

### Cytoplasmic content is inappropriately exchanged between *mos^-/-^* eggs and polar bodies

To test our hypothesis that *mos^-/-^* eggs maintain a connection between the egg and the polar body, we asked if components within the egg cytoplasm could pass into the polar body. To this end, we injected wildtype and *mos^-/-^* eggs with dextran conjugated to a fluorescent dye and then monitored whether it could move into the polar body. We first injected the eggs with dextran at 13.5 hours after prophase I release (Figure 3A). We chose this timepoint because polar body divisions in *mos^-/-^*eggs occur 13-15 hours after prophase I release. In wildtype eggs, we observed that the fluorescent dextran did not pass into the polar body (Figure 3B). In contrast, 50% of *mos^-/-^* eggs had fluorescent dextran that passed from the egg into the polar bodies, suggesting that cytoplasmic content can pass between the two compartments (Figure 3C-D). Interestingly, the percentage of *mos^-/-^* eggs that had fluorescent dextran in the polar body is similar to the percentage that assembled bipolar spindles in the polar body, suggesting that the eggs that have cytoplasmic connections can assemble spindles in the polar body (Figure 1E, 3D).

**Figure 3.**
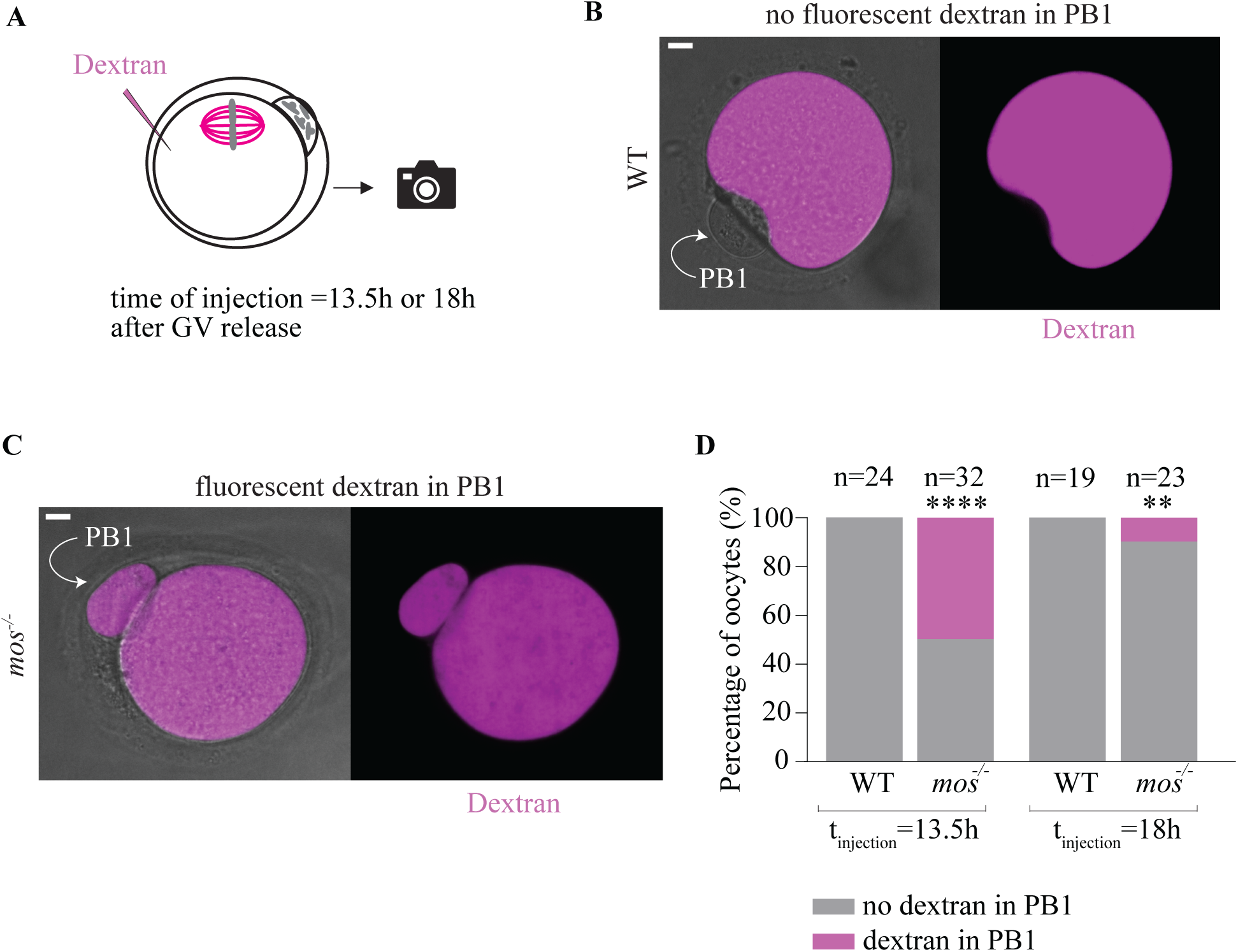
*mos^-/-^* eggs have a prolonged exchange of cytoplasm between the egg and polar body. **(A)** Schematic of experiment outline. Wildtype and *mos^-/-^* eggs were injected with dextran at 13.h and 18h after prophase I release and subsequently imaged. **(B)** Representative images of a wildtype injected at 13.5h after prophase I release. **(C)** Representative images of a *mos^-/-^* egg injected at 13.5h after prophase I release. **(D)** Percentage of wildtype and *mos^-/-^* eggs that show no dextran or dextran in the polar body with dextran injection at 13.5h or 18h after prophase I release (n indicates the number of eggs injected and imaged per condition at each timepoint; ****, P < 0.0001, **, P = 0.068 two-tailed Fisher’s exact test; 3 or more independent experiments were conducted per condition) GV, germinal vesicle; WT, wildtype; PB, polar body.

We then asked if the cytoplasm of the egg and the polar body remained connected or had a delayed separation. We injected fluorescent dextran into wildtype and *mos^-/-^* eggs 18 hours after prophase I release. As expected, we did not observe fluorescent dextran in the polar body of wildtype eggs (Figure 3D). The percentage of *mos^-/-^* eggs with fluorescent dextran passing from egg to polar body reduced to approximately 10% when dextran was injected 18 hours after prophase I release. Therefore, *mos^-/-^* eggs and polar bodies eventually separated into two compartments. These results suggest that there is a delayed separation between *mos^-/-^* eggs and polar bodies, and that cell cycle regulators can pass into the polar body to allow spindle assembly and polar body division with similar timing as anaphase II onset in the *mos^-/-^* eggs that activate parthenogenetically.

### The meiotic midbody is defective in *mos^-/-^* eggs

Given the movement of dextran from the polar body to the egg, we next asked whether cytokinesis defects contribute to the delayed separation of egg and polar body cytoplasm in *mos^-/-^* mutants. From our time-lapse imaging, the membrane ingresses between the egg and polar body, suggesting actomyosin ring assembly and contraction. We therefore assessed the assembly of the meiotic midbody (mMB), a transient organelle that forms during early to late telophase I as the egg and polar body separate. At the mMB, the microtubules have a unique orientation and mMB substructure made up of Centralspindlin complex assemblies (Hu et al., 2012; Jung et al., 2023). In somatic cells, the Centralspindlin component MKLP1 (mitotic kinesin-like protein 1) forms a ring at the midbody, whereas oocytes form both a ring and a cap, with the cap pointing into the polar body. The mMB cap creates a barrier to prevent nascent proteins from entering the polar body, ensuring their retention within the egg (Jung et al., 2023).

We asked if the mMB structure assembled abnormally in *mos^-/-^*oocytes. We used immunofluorescence and super-resolution microscopy to observe MKLP1 localization in fixed telophase I-staged oocytes. The majority of wildtype oocytes had the ring and cap substructure, as previously observed (Figure 4A) (Jung et al., 2023). In contrast, approximately 90% of *mos^-/-^* oocytes had a defective substructure (Figure 4B-E). Most of the *mos^-/-^* oocytes had an MKLP1 ring, but the cap was absent (Figure 4B). We also observed substructures with a ring and a faint cap that pointed into the oocyte instead of the polar body, or an entirely abnormal structure, appearing fluffy instead of well-defined (Figure 4C-D). These results indicate that *mos^-/-^* oocytes have a defective mMB substructure.

**Figure 4.**
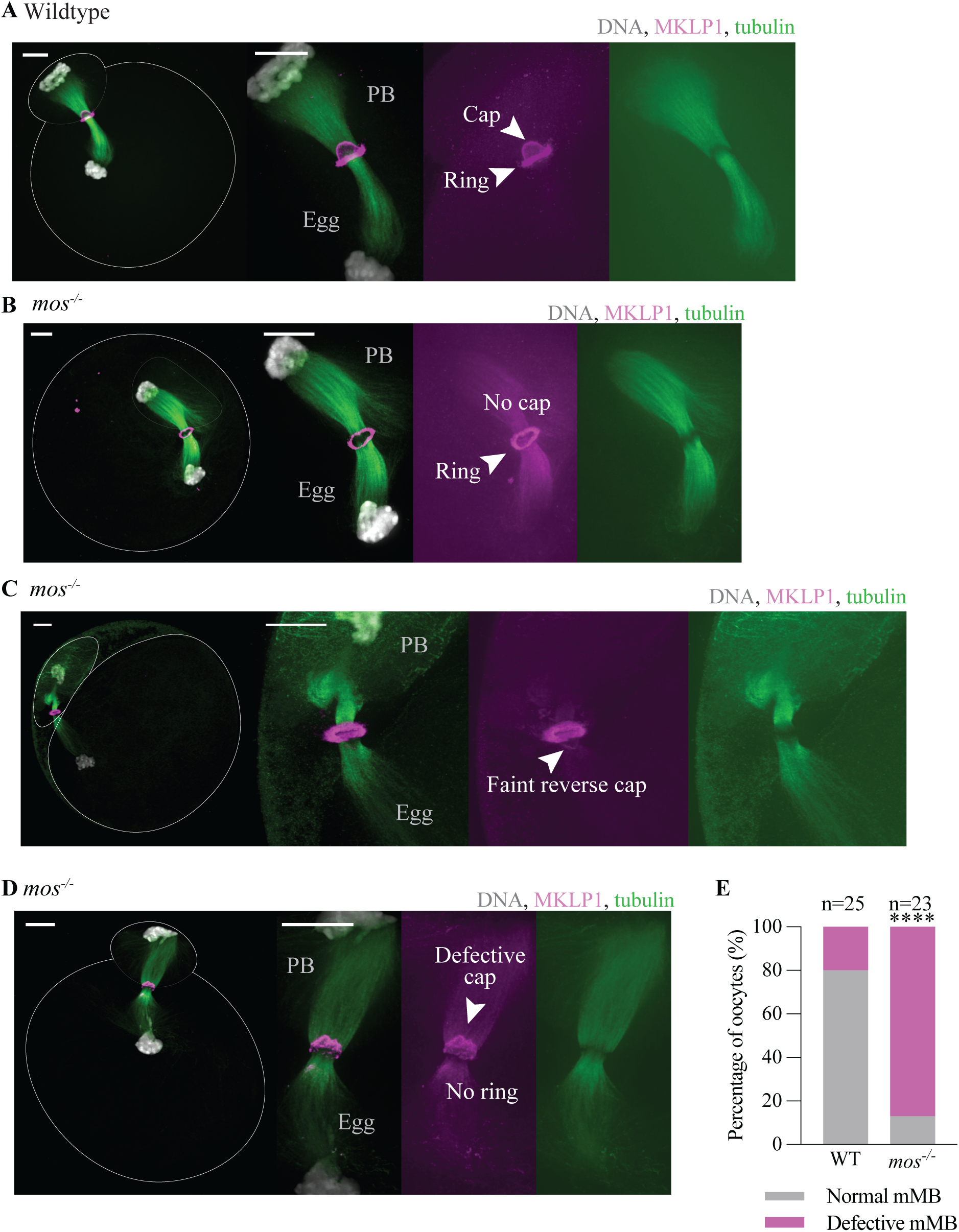
*mos^-/-^*eggs have defects in meiotic midbody formation. **(A-D)** Representative immunofluorescence images of wildtype (A) and *mos^-/-^* eggs (B-D) showing localization of MKLP1 (magenta), tubulin (green), and DNA (white). Arrowheads indicate structure features of MKLP1. Wildtype and *mos^-/-^* oocytes were fixed 9h-11h30min after prophase I release. Scale bars, 10µm. **(E)** Percentage of wildtype and *mos^-/-^* eggs with normal or defective MKLP1 structure (n indicates the number of eggs imaged per condition; 3 or more independent experiments conducted per condition; ****, P < 0.0001, two-tailed Fisher’s exact test).

We next examined the localization of a component of the core mMB, PRC1 (protein regulator of cytokinesis 1) (Hu et al., 2012; Jung et al., 2023). In most wildtype eggs, PRC1 localized as two disk-like structures at telophase I with a dome-like disk pointing into the polar body (Figure S3A)(Jung et al., 2023). In 65% of *mos^-/-^*oocytes, we observed that PRC1 localization was defective, showing either absence of one disk, or a defective structure (Figure S3B-E). Overall, our results demonstrate that *mos^-/-^* oocytes have defective mMB structures. Therefore, our findings support the model that defects in the mMB allows the passage of cytoplasmic content between the egg and the polar body, allowing spindle assembly and chromosome segregation in the polar body with similar timing as anaphase II onset in the *mos^-/-^* eggs that parthenogenetically activate.

### Polar bodies in *mos^-/-^* eggs enter a new cell cycle and undergo DNA replication

Altogether, our results suggest that the egg and the polar body maintain prolonged connection that allows the polar body to undergo an additional round of chromosome segregation. Our next question was whether the polar body could initiate a new cell cycle and replicate its chromosomes, possibly contributing to the cell cycle divisions of the tumor. To address this question, we used a 5-ethynyl-2’-deoxyuridine (EdU) clickIT assay, which allows the detection of newly synthesized DNA through incorporation of EdU, a thymidine analog (Salic and Mitchison, 2008). We matured wildtype and *mos^-/-^*eggs in EdU-containing media and fixed them 38-40 hours after prophase I release (Figure 5A). We then incubated with the Click-IT reaction mix and stained for DNA with DAPI (4,6-Diamidino-2-Phenylindole, Dihydrochloride). We imaged the EdU to assess new DNA synthesis and DAPI to visualize the DNA. As expected for eggs arrested at metaphase II, we never observed EdU staining in wildtype eggs (Figure 5B). In *mos^-/-^* mutants, we observed EdU staining in both eggs and polar bodies with decondensed chromatin, suggesting that DNA replication could occur in both the egg and the polar body (Figure 5C-E). It is important to note that this experiment is a snapshot, not a time-lapse, showing those eggs and polar bodies that were actively undergoing DNA replication. Therefore, there are likely more *mos^-/-^* eggs and polar bodies that undergo DNA replication at other points. However, our results show that in *mos^-/-^* mutants, both eggs and polar bodies can initiate a new cell cycle and undergo DNA replication. Therefore, these data support the model that polar bodies contribute to abnormal mitotic divisions initiated in *mos^-/-^* eggs that escaped meiosis.

**Figure 5.**
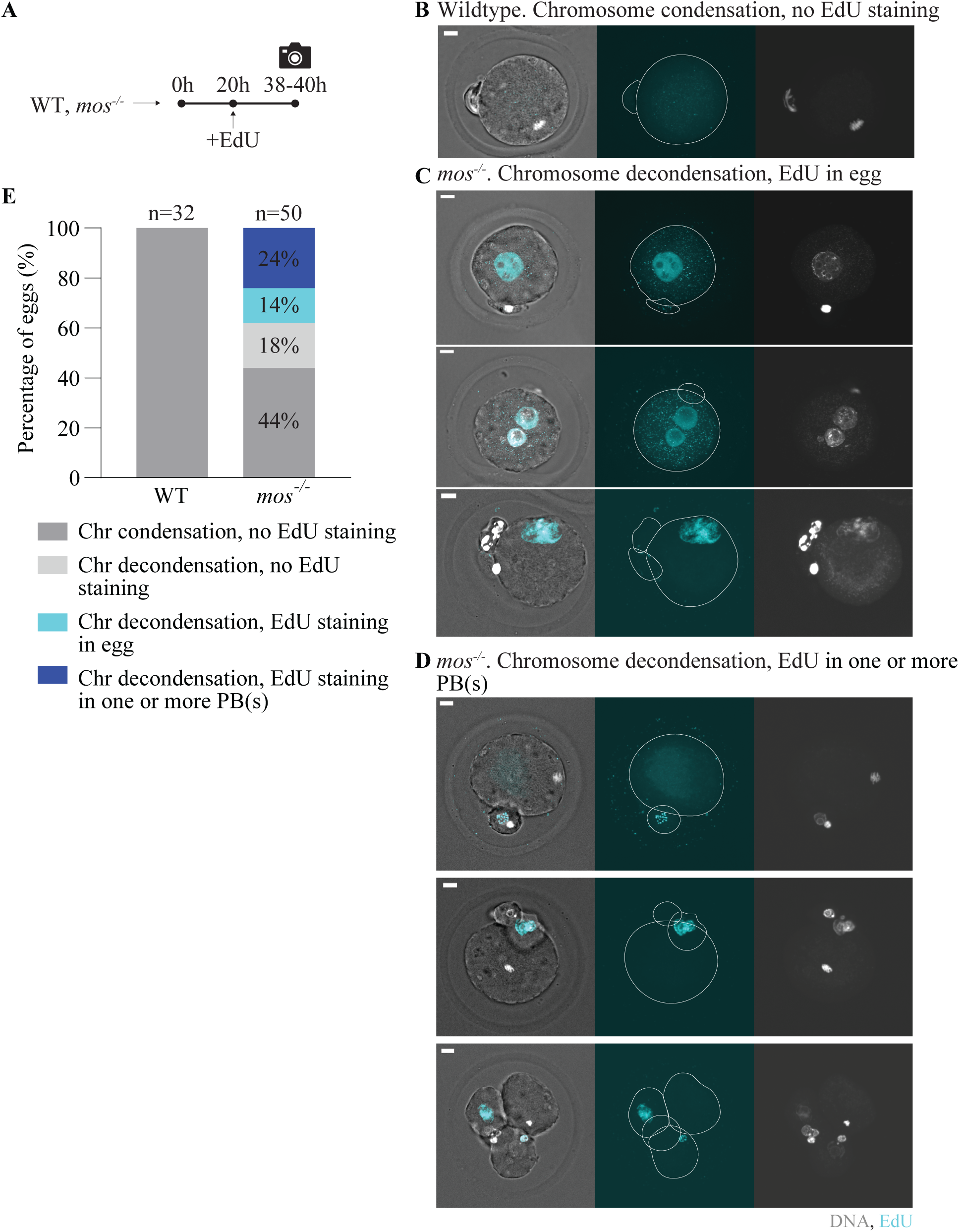
DNA replication occurs in *mos^-/-^*eggs and polar bodies. **(A)** Schematic of experiment outline. Wildtype and *mos^-/-^* eggs were incubated in EdU-containing media and fixed 38-40h after prophase I release. Eggs were incubated with the Click-IT reaction mix and immunostained for DNA. **(B-D)** Representative immunofluorescence images of wildtype (B) and *mos^-/-^* eggs (C-D) with DNA (white) and EdU staining (cyan). Scale bars, 10µm. **(E)** Percentage of wildtype and *mos^-/-^* eggs exhibiting condensed or decondensed chromosome as well as the presence of EdU staining (n indicates the number of eggs imaged per condition; 3 or more independent experiments conducted per condition). WT, wildtype. EDU, 5-ethynyl-2’-deoxyuridine.

### Polar bodies in *mos^-/-^* eggs can survive and segregate chromosomes after separation from the egg

We then asked if the first extruded polar bodies from *mos^-/-^* eggs could undergo abnormal divisions when separated from the egg and zona pellucida. We matured wildtype and *mos^-/-^*oocytes and dissolved their zona pellucida and then allowed the first extruded polar body (PB1) to separate from the egg. We then collected the polar bodies that had separated from the eggs and either monitored morphological changes every hour or used time-lapse imaging to monitor the chromatin and spindle with SirDNA and SPY-555 tubulin, respectively (Figure 6A-D). 85% of isolated wildtype polar bodies typically survived for less than 3 hours and did not divide (Figure 6A, D). The remaining 15% survived between 3-8 hours and did not divide. In contrast, *mos^-/-^* polar bodies typically survived longer, with approximately 50% undergoing chromosome segregation and most of those surviving for more than 8 hours (Figure 6B-D). Interestingly, the isolated *mos^-/-^* polar bodies also underwent rapid morphological changes with incomplete cytokinesis (Figure 6B-C, S4A). These changes were not typical to the polar bodies that remained connected to the egg and with the zona pellucida intact. Therefore, the egg and zona pellucida may provide connections that stabilize the morphology of the polar body in *mos^-/-^* eggs that undergo additional divisions. In conclusion, these experiments show that the polar bodies isolated from *mos^-/-^* eggs can survive and undergo chromosome segregation, and attempt cell division, supporting the ability to contribute to germ cell tumor formation in vivo.

**Figure 6.**
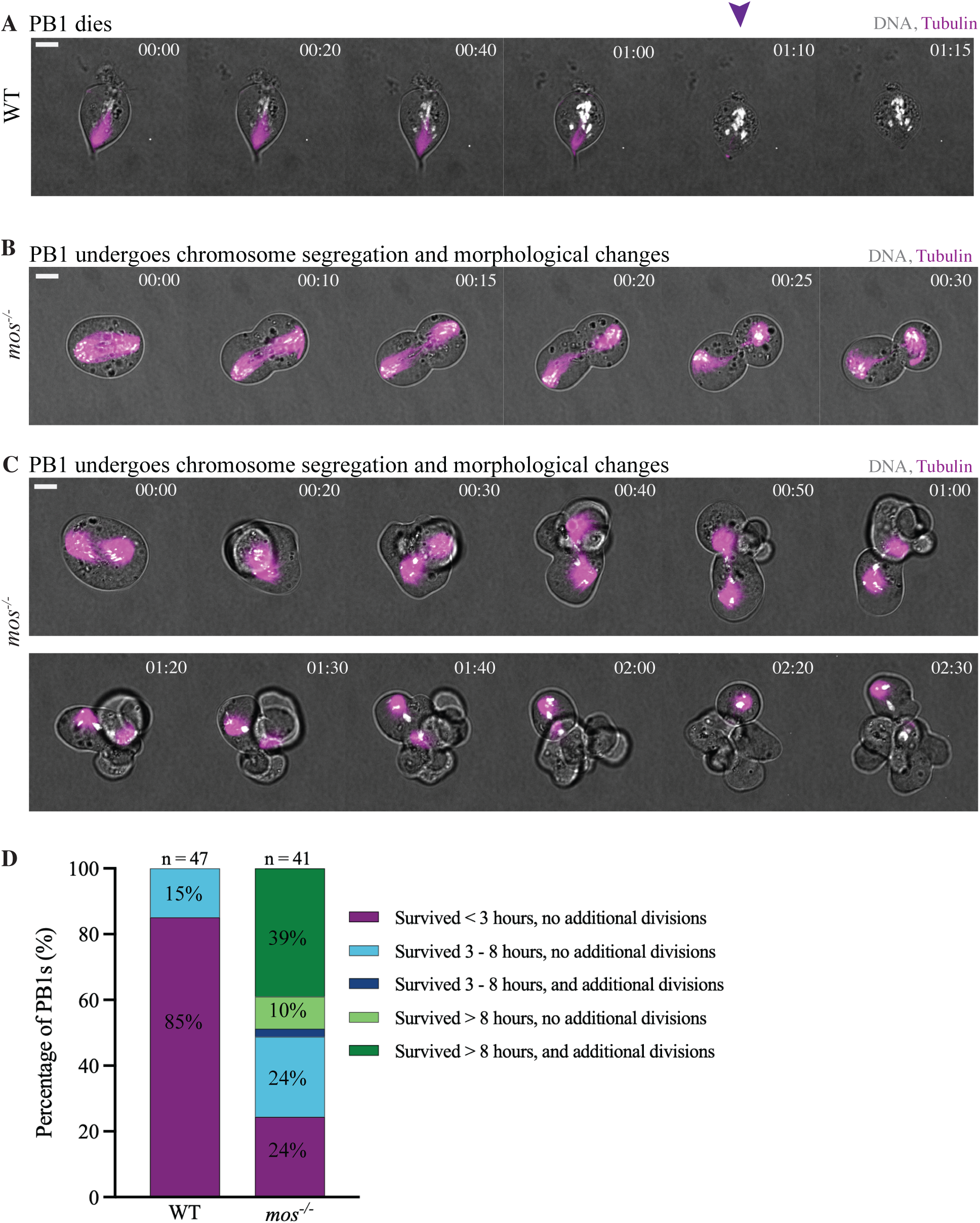
Isolated and cultured *mos^-/-^* polar bodies can survive and undergo chromosome segregation. **(A-C)** Representative time-lapse images of wildtype (A) and *mos^-/-^* (B-C) isolated polar bodies. SPY555tubulin and SirDNA were added to visualize tubulin (magenta) and DNA (white), respectively. Purple arrowhead shows the timepoint the wildtype polar body dies. Scale bars, 10µm. Time 0 represents the time at which the movie was initiated. **(D)** Percentage of wildtype and *mos^-/-^* isolated polar bodies that survive and undergo additional divisions. n indicates the number of isolated polar bodies analyzed per condition. 2 or more independent experiments were conducted per genotype. GV, germinal vesicle; WT, wildtype; PB1, first extruded polar body.

## DISCUSSION

MOS/MAPK signaling plays critical roles in several aspects of oocyte meiosis, including cell cycle progression, normal spindle assembly, filamentous-actin-mediated cortical remodeling, and RNA regulation (Cao et al., 2020; Chaigne et al., 2013; Chaigne et al., 2015; Choi et al., 1996; Deng et al., 2005; Kalous et al., 2018; Lefebvre et al., 2002; Sha et al., 2017; Terret et al., 2003; Verlhac et al., 1996; Verlhac et al., 2000b). Among its important functions, MOS is pivotal for maintaining the metaphase II arrest (Dupré et al., 2011; Hörmanseder et al., 2013). In the absence of MOS, a subset of eggs exits metaphase II and proceeds through aberrant, inappropriate divisions (Colledge et al., 1994; Hashimoto et al., 1994). Notably, approximately one-third of *mos^-/-^* female mice develop germ cell tumors, suggesting that some of these aberrantly dividing eggs fail to ovulate and instead undergo uncontrolled divisions within the ovary.

We used time-lapse imaging to gain further insight into the additional divisions in *mos^-/-^* eggs. We found that the first polar body assembles bipolar spindles and segregates chromosomes with similar timing to anaphase II onset in the *mos^-/-^*eggs that undergo parthenogenetic activation (Figure 1). Because *mos^-/-^* eggs are known to have larger polar bodies (Choi et al., 1996; Verlhac et al., 2000a), a potential mechanism could have been that they took up more cytoplasm and therefore more cell cycle regulators, leading to a continuation of the cell cycle in the polar body. However, we found that the size of the polar body did not correlate with spindle assembly. Both larger and normal-sized polar bodies assembled spindles and divided their chromosomes (Figure 2). Our results showed that the major predictor of spindle assembly in the polar body was not polar body size, but instead whether the egg underwent anaphase II; most activated eggs display bipolar spindles in the polar body (Figure 1E).

The finding that spindle assembly and chromosome segregation take place in the first polar bodies of *mos^-/-^* mutants was surprising because wildtype polar bodies are typically non-functional cells that will undergo an apoptotic-like degeneration (Miao et al., 2004; Wakayama and Yanagimachi, 1998). However, in *mos^-/-^*eggs, polar bodies were less likely to degenerate and instead 53% formed bipolar spindles and divided (Figure 1, S1)(Choi et al., 1996). Although the polar body divisions mostly occurred in *mos^-/-^*eggs that underwent parthenogenesis (Figure 1B), the phenotype is not a feature of parthenogenesis. Wildtype eggs that were chemically activated to undergo parthenogenesis did not have polar body divisions, suggesting that the loss of MOS is causing the polar body divisions.

We propose a model that the prolonged connection between the *mos^-/-^*egg and polar body prevented polar body degeneration by allowing the cytoplasmic contents to pass between the two spaces. Our experiments showed that *mos^-/-^* but not wildtype eggs could pass injected dextran from the egg into the polar body at a timepoint when spindles typically assembled in the polar body (Figure 3). The trigger that promotes polar body degeneration is not known; it could be a signal that promotes degeneration or the lack of a proliferation or growth signal that promotes degeneration. However, the exchange of cytoplasmic content between the egg and the polar body could potentially dilute the degeneration signal or allow a proliferation signal to enter the polar body for spindle assembly and chromosome segregation.

The exchange of cytoplasmic content between the *mos^-/-^*egg and polar body is likely due to a disrupted mMB structure. In somatic cells, the midbody component MKLP1 forms a ring structure (Hu et al., 2012). In oocytes, MKLP1 forms a ring and a cap, with the cap pointing towards the polar body (Jung et al., 2023). The midbody is an active site for new translation in both somatic cells and oocytes and the cap is thought to prevent nascent polypeptides from entering the polar body. Experiments disrupting the cap showed leakage of the nascent polypeptides into the polar body. We find that the mMB structure is defective in *mos^-/-^*eggs, most presenting with an MKLP1 ring, but lacking the cap (Figure 4). In the future, it will be interesting to determine how MOS/MAPK signaling regulates mMB assembly. We note that there is a prolonged intracellular bridge connection between the two spaces, but at a later timepoint, fluorescent dextran was unable to pass into the polar body. Therefore, abscission does eventually occur in *mos^-/-^* eggs, but at timepoints after spindle assembly within the polar body.

Overall, our results support the model that defective meiotic midbody cap permits cytoplasmic exchange between the egg and the polar body. We propose that two key factors contribute to the aberrant polar body divisions observed in *mos^-/-^* eggs: i) parthenogenetic activation of the egg, and ii) a prolonged intracellular bridge between the egg and the polar body that enables leakage of cell cycle regulators from the egg into the polar body. Another key finding from this work is that, following the first meiotic division, polar bodies can enter a new cell cycle, as evidenced by the initiation of DNA replication (Figure 5). These polar bodies can be isolated and survive in culture, segregate chromosomes, and undergo morphological changes (Figure 6). These results indicate that polar bodies can undergo abnormal mitotic divisions, which may ultimately contribute to ovarian germ cell tumor formation.

Our results also have implications for understanding human infertility. Variants of the *MOS* gene have been identified in women undergoing fertility treatment (Jiao et al., 2022; Zhang et al., 2021; Zhang et al., 2022). *In vitro* fertilization of the eggs from these individuals often results in abnormal early embryonic divisions, resulting in developmental arrest and fragmentation. Embryos derived from these eggs have varying cell sizes. Inspired by our results, it would be interesting to determine if polar bodies themselves contribute to these abnormal divisions in human eggs and embryos.

## METHODS

### Mouse strains and genotyping

All animal procedures were conducted at the Geisel School of Medicine at Dartmouth College in accordance with protocols approved by the Institutional Animal Use and Care Committee at Dartmouth in accordance with NIH guidelines (IACUC Protocol# 00002303). Sexually mature wildtype and *mos^-/-^* knockout mice, aged 8-12 weeks, were used for all experiments. The originating *mos^+/-^*heterozygous female and male mice were obtained from the Jackson Laboratories (Cat# 002723). The knockout mutation was made by introducing a neomycin resistant cassette upstream of the *Mos* kinase domain in a C57BL/6J background. Experimental animals were generated by mating heterozygous females to either homozygous or heterozygous males. Genotyping was performed using a common forward primer (5’-TCAGCTGCAGAGAACAACTGA-3’) and two allele-specific reverse primers to either detect the wildtype *Mos* (5’-GTGTACGTGCCCCCTATGTG-3’), or the neomycin resistance cassette in the *Mos* gene (5’-GCCAGAGGCCACTTGTGTAG-3’). Mice were housed under standard conditions, including a 12-hour dark/light cycle, ambient temperature (72+/-3°F), 30-70% humidity, and access to food and water *ad libitum*.

### Mouse oocyte isolation and maturation

Prior to each experiment, female mice were injected intraperitoneally with 5 I.U. of pregnant mare serum gonadotropin (PMSG; BioVendor R&D RP1782725000) 48h before oocyte collection. Prophase I-arrested oocytes were collected from ovaries and placed in minimal essential medium (MEM) prepared with Earle’s salts (Sigma M0268), 0.1g pyruvate (Sigma P4562), 1mL Gentamycin (Sigma G1272), 25mL 1M HEPES pH 7.3 (Sigma H3784), 3g PVP pH 7.3 (Sigma P2307). To prevent spontaneous meiotic resumption, 2.5μM milrinone (Sigma-Aldrich, M4659) was added. Oocytes were then cultured in Chatot, Ziomek, and Bavister (CZB) media containing 81.6mM NaCl (Sigma S5886), 4.8mM KCl (Sigma P5405), 1.2 mM KH_2_PO_4_ (Sigma P5655), 1.2 mM MgSO_4_ 7H_2_O (Sigma M7774), 0.27 mM Pyruvic acid (Sigma P4562), 1.7 mM CaCl_2_ 2H_2_O (Sigma C7902), 30.8 mM DL-Lactic acid (Sigma L7900), 7mM Taurine (Sigma T0625), 0.1mM EDTA (Sigma E5134), 25mM NaHCO_3_ (Sigma S5761), 0.1% Gentamicin, (Sigma G1272), 0.3% BSA (Sigma A4503). Oocytes were cultured in CZB without milrinone to allow meiotic progression. All cultured oocytes were maintained in a humidified incubator at 37°C with 5% CO_2_. Each experiment was independently performed at least twice, with multiple control and experimental animals included in each replicate.

To induce parthenogenetic activation, wildtype eggs were matured in CZB medium for 15-16h following prophase I release. Mature eggs were then washed and incubated in Ca^2+^-free CZB medium with 10mM strontium chloride (Sigma Aldrich 25521) for 2h. Successful activation was assessed by the appearance of a pronucleus and/or extrusion of a second polar body. Activated eggs were subsequently washed and maintained in CZB for imaging.

### Live imaging of mouse oocytes and eggs

For live imaging, SirDNA and SPY555-tubulin (Spirochrome) were reconstituted in DMSO and added to CZB medium at final concentrations of 100-250nM, 45min to 1h prior to imaging. Live imaging was performed using a CSU-SoRa spinning disk confocal microscope, equipped with a humidified chamber set to 5% CO_2_ and 37°C. Oocytes and eggs were imaged in 250μL of CZB medium placed in chambered cover glass dishes (Cellvis C18-1.5H) using a 40x Plan Apochromat λS 1.15 numerical aperture water-immersion objective. All experimental repetitions were imaged on the same microscope under identical imaging settings. Tubulin and DNA signals were captured using exposure times of 200ms and 30ms, respectively, with neutral-density filters transmitting 25% (tubulin) and 5% (DNA) of the excitation light. For time-lapse imaging, Z stacks of 13 optical sections at 4μm intervals were acquired every 20min for 24h, allowing for high-resolution monitoring of meiotic progression.

### EdU Click-IT assay

To assess DNA replication, incorporated EdU was detected using the Click-iT EdU Alexa Fluor 647 Imaging Kit (Invitrogen C10340) following the manufacturer’s instructions. Wildtype and *mos^-/-^* eggs were first cultured in CZB medium for 20h after prophase I release. Eggs were then washed and incubated in CZB supplemented with 10μM EdU for 12-14h, followed by fixation in 3.7% paraformaldehyde for 15min at room temperature. After fixation, eggs were washed twice in 3% BSA in PBS, then permeabilized with 0.5% Triton X-100 for 15min at room temperature. Following two additional washes with 3% BSA in PBS, eggs were incubated with the Click-iT reaction cocktail for 30min at room temperature, protected from light. Eggs were then washed once with 3% BSA in PBS and then mounted on a slide containing 5μL of Vectashield mounting medium (Vector Laboratories, H-1000) containing DAPI (Life Technologies D1306) at a 1:170 dilution. Imaging was performed using a CSU-SoRa spinning disk confocal microscope equipped with a 60x Plan Apochromat λD 1.42 numerical aperture oil-immersion objective. EdU and DNA were visualized using exposure times of 100ms and 700ms, respectively, with neutral-density filters that transmitted 25% of the excitation light for both channels. Z-stacks were acquired at 0.4μm intervals.

### Dextran microinjections

To assess separation of egg and polar body, wildtype and *mos^-/-^*eggs were injected with 10mg/mL Texas Red^TM^-labeled fluorescent dextran (Invitrogen D3328) either 13.5h or 18h following prophase I release. Dextran was reconstituted in cell culture-grade water (Corning 25-055-CM) and stored at -20°C in the dark until use. Injected eggs were imaged 5-10min post-injection using a CSU-SoRa spinning disk confocal microscope equipped with a humidified incubation chamber set to 5% CO_2_ and 37°C. Imaging was performed in 250μL of CZB medium placed in chambered cover glass dishes (Cellvis C18-1.5H) using a 40x Plan Apochromat λS 1.15 numerical aperture water-immersion objective. Dextran fluorescence was visualized using an exposure time of 100ms with neutral-density filters transmitting 20% of the excitation light. An image was acquired on the plane of focus containing the polar body.

### Immunofluorescence

For immunostaining of DNA and tubulin, wildtype eggs and *mos^-/-^* eggs were fixed 24h after prophase I release in 2% paraformaldehyde prepared in phosphate-buffered saline (PBS) for 20min at room temperature. Following fixation, eggs were transferred to blocking solution consisting of 0.3% BSA, 0.01% Tween-20, 0.02% NaN_3_ in PBS and stored at 4 °C for no longer than 48h. Eggs were then permeabilized in 0.3% BSA, 0.1% Triton X-100, 0.02% NaN_3_ in PBS for 20min and washed 3 times in blocking solution, each for 10min. For tubulin detection, eggs were incubated for 1hr at room temperature in blocking solution containing Alexa Fluor 488 conjugated anti-alpha-tubulin rabbit monoclonal antibody (11H10; Cell Signaling Technology 5063S; dilution of 1:100) in a humidified chamber. Following incubation, eggs were washed three times for 10mins each in blocking solution. Eggs were then mounted in 5μL of Vectashield mounting medium (Vector Laboratories H-1000) containing DAPI (Life Technologies D1306; 1:170) to detect DNA. Images were acquired with a CSU-SoRa spinning disk confocal microscope with a 60x Plan Apochromat λD 1.42 numerical aperture oil-immersion objective. Tubulin and DNA were imaged with exposure times of 700-900ms, using neutral-density filters transmitting 20-25% of the excitation light. Z-stacks were collected at 0.3μm intervals.

To visualize midbody components MKLP1 and PRC1, along with tubulin and DNA, wildtype and *mos^-/-^* eggs were fixed using 2% paraformaldehyde 9h-11.5h post prophase I release. Fixed eggs were permeabilized for 20min at room temperature, followed by three 10min washes in blocking solution. Eggs were then incubated for 1hr at room temperature in a humidified chamber with primary antibody against either MKLP1 or PRC1, diluted in blocking solution. The primary antibodies used were rabbit anti-MKLP1 polyclonal antibody (Novus NBP2-56923; 1:100) and rabbit anti-PRC1 polyclonal antibody (Proteintech 15617-1-AP; 1:100). After primary incubation, eggs were washed three times, 10 minutes each, in blocking solution. Next, eggs were incubated in blocking solution with Alexa Fluor488-conjugated mouse anti-alpha-tubulin monoclonal antibody (DM1A; Cell Signaling Technology 8058S; 1:100) and the appropriate secondary antibody to detect either MKLP1 or PRC1: Alexa Fluor 568 donkey anti-rabbit IgG (H+L; Life Technologies A10042; 1:200) or Alexa Fluor 568 donkey anti-rabbit IgG (H+L; Invitrogen, 1:200), respectively. Following incubation, the eggs were then washed three times with blocking solution for 10 minutes each and mounted in 5μL of Vectashield mounting medium (Vector Laboratories H-1000) containing DAPI (Life Technologies D1306; 1:170) to detect DNA.

Oocyte images were acquired with a CSU-SoRa spinning disk confocal microscope equipped with a 60x Plan Apochromat λD 1.42 numerical aperture oil-immersion objective. MKLP1 or PRC1, tubulin, and DNA were visualized with exposure times of 800 (MKLP1/PRC1), 800 (tubulin) and 600ms (DAPI), with neutral-density filters transmitting 25%, 25%, and 20% of excitation light, respectively. Z-stacks were acquired at 0.4μm intervals, covering a total depth of 30-35μm. High-resolution images of MKLP1 or PRC1, tubulin, and DNA were captured using the super-resolution mode of the CSU-SoRa system with 2x SoRa magnification and the same objective. For these images, MKLP1 or PRC1, tubulin, and DNA were visualized with exposure times of 400ms, 900ms, and 600ms, respectively with all channels using 20% neutral-density filters. Z-stacks were collected at 0.2μm intervals across a total depth of 30-35μm.

### First polar body (PB1) cultures

Oocytes were harvested and released from prophase I in CZB medium at 37°C with 5% CO₂. After 3 hours, oocytes that had undergone germinal vesicle breakdown (GVBD), identified by the absence of a pronucleus, were isolated and then denuded, as previously described (Fan et al., 2021). Briefly, oocytes were bathed through droplets of acidic Tyrode’s solution (Sigma Aldrich MR-004-D) for 40 seconds, followed by 20 seconds in CZB containing 0.05% Proteinase K (ThermoFisher, EO0491). Denuded oocytes were washed in fresh CZB droplets and then incubated in CZB at 37°C with 5% CO₂. After 10 hours, cultures were monitored for abscised first polar bodies (PB1s), which appeared as free-floating cells in the medium. Naturally abscised PB1s were transferred to fresh CZB culture medium and maintained at 37°C with 5% CO₂. Separately cultured PB1s were examined hourly under a stereoscope to assess viability and cell division-like morphology.

To further analyze additional-division-like phenotype observed for *mos^-/-^* PB1s, WT and *mos^-/-^* PB1s were subjected to live-cell imaging (described above) in the presence of 250nM SiR-DNA and SPY555-tubulin (Spirochrome). Images were acquired every 5 minutes for 12 hours. For Z-series acquisition, a 4 µm step size was used to cover a total depth of 60 µm.

## ACKNOWLEDGEMENTS

We thank the Lacefield Lab for reading the manuscript and Binyam Mogessie for valuable discussions. We thank our core facilities through Dartmouth Mouse Modeling Shared Resource (RRID SCR – 021284) which is supported by Dartmouth Cancer Center Support Grant 5P30CA023108 from the NCI. SL is supported by the bioMT through NIH NIGMS grant P20-GM113132. KS is supported by R35-GM136340. Use of the Nikon SoRa spinning disk confocal at the Life Sciences Light Microscopy facility at Dartmouth was supported through NIH award S10OD032310 to Dr. Yashi Ahmed.

## AUTHOR CONTRIBUTIONS

**Figure S1.**
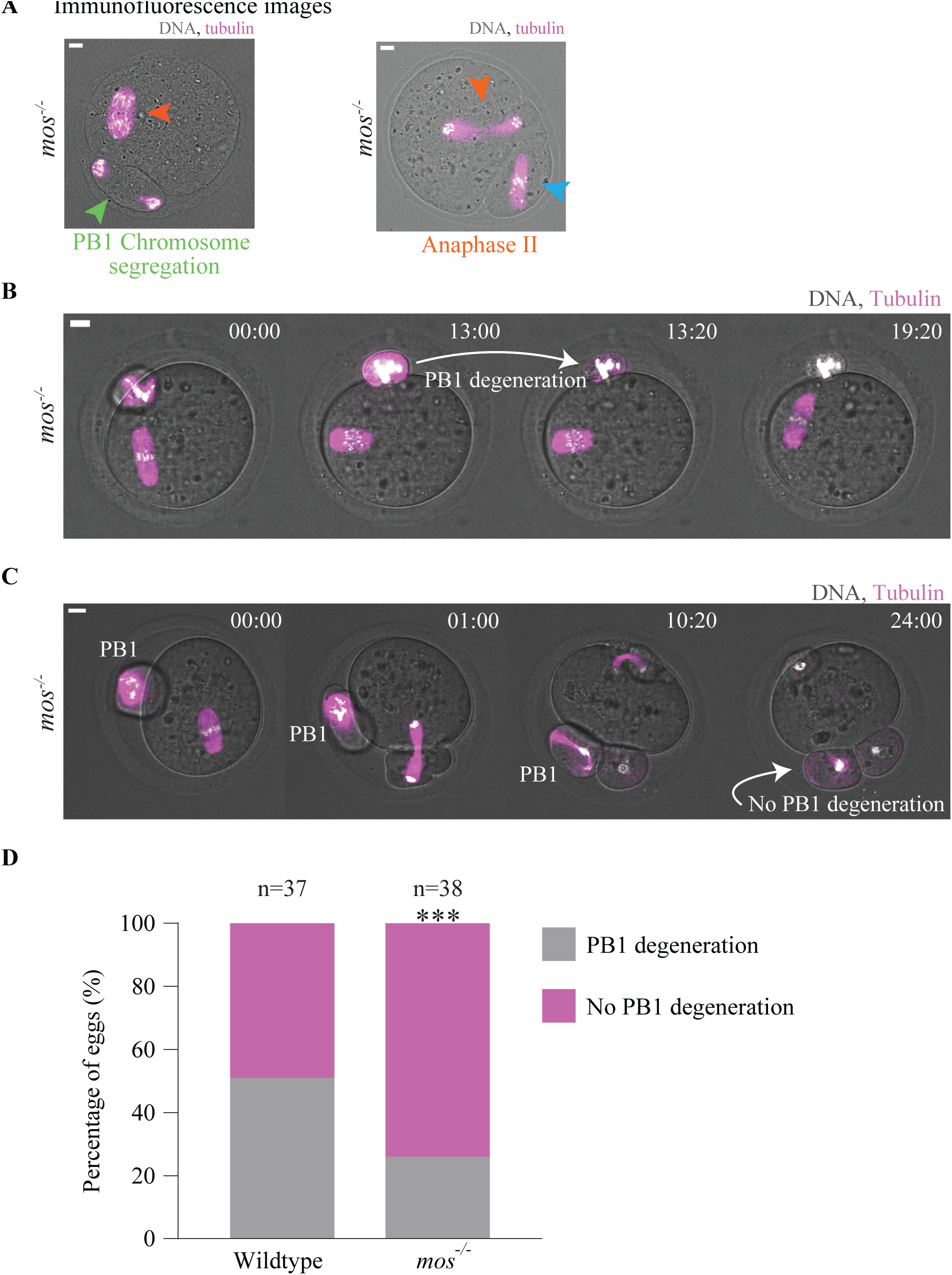
Fewer polar bodies degenerated in *mos^-/-^* eggs. **(A)** Representative immunofluorescence images of *mos^-/-^* eggs showing localization of DNA (white) and tubulin (magenta). Blue arrowhead indicates bipolar spindle formation inside the PB; orange arrowheads indicate anaphase II in the egg; green arrowhead indicates chromosome segregation inside the PB. Scale bars, 10µm. **(B-C)** Time-lapse images of *mos^-/-^*eggs (B-C) imaged starting 13-15h after prophase I release for 24 hours. Polar body degeneration is highlighted. Scale bars, 10µm. **(D)** Percentage of wildtype and *mos^-/-^* eggs with degenerated polar bodies during the time of the imaging. (n indicates the number of eggs imaged per condition; 3 or more independent experiments conducted per condition; ****, P < 0.0001, two-tailed Fisher’s exact test). GV, germinal vesicle; WT, wildtype; PB, polar body.

**Figure S2.**
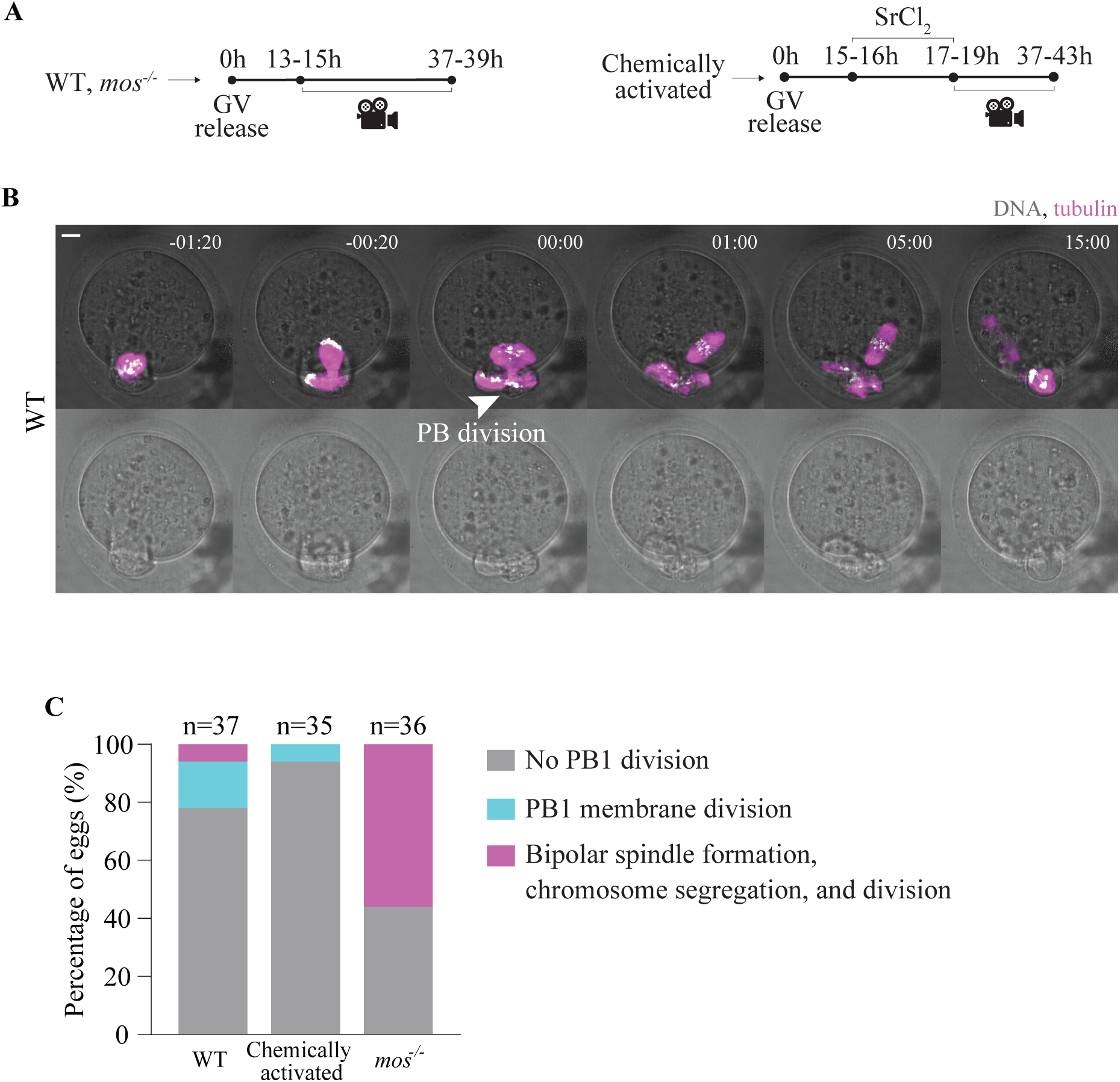
A subset of wildtype eggs has abnormal polar body membrane division without prior bipolar spindle formation after prolonged metaphase II arrest. **(A)**. Schematic of experiment outline. **(B)** Representative time-lapses of a wildtype egg (B) undergoing PB membrane division. SPY555tubulin and SirDNA were added to image tubulin (magenta) and DNA (white), respectively. Time 0 represents the moment of PB membrane division. Arrowhead indicates PB membrane division. Scale bars, 10µm. **(C)** Percentage of wildtype, chemically activated, and *mos^-/-^* eggs undergoing PB membrane division (n indicates the number of eggs imaged per condition; 3 or more independent experiments conducted per condition; GV, germinal vesicle; WT, wildtype; PB, polar body).

**Figure S3.**
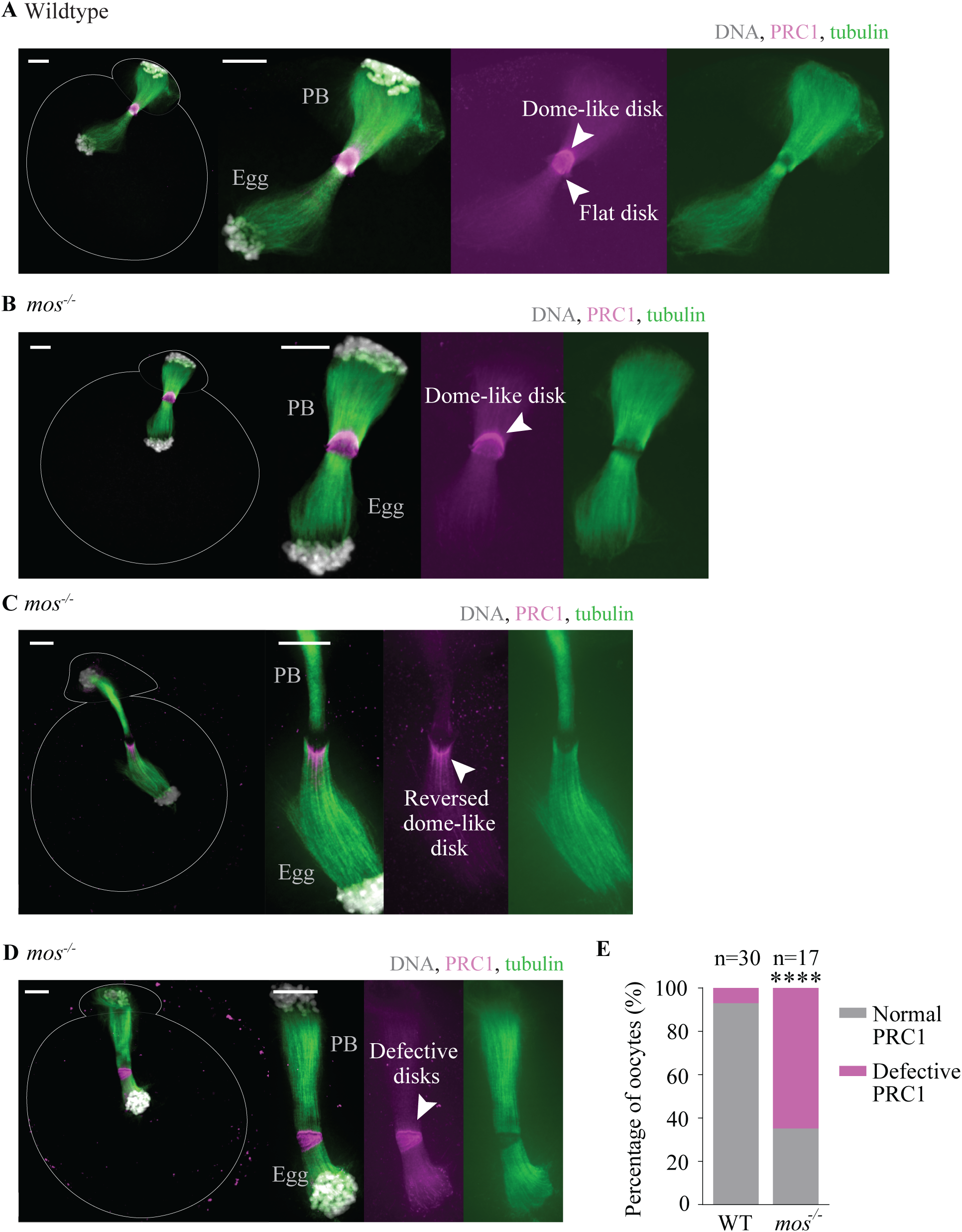
(A-D) *mos^-/-^*eggs exhibit defects in PRC1 localization. Representative immunofluorescence images of wildtype (A) and *mos^-/-^* eggs (B-D) showing localization of PRC1 (magenta), tubulin (green), and DNA (white). Arrowheads indicate structure features of PRC1. Wildtype and *mos^-/-^* oocytes were fixed 9h-11h30min after prophase I release. Scale bars, 10µm. **(E)** Percentage of wildtype and *mos^-/-^*eggs exhibiting a normal or defective PRC1 structure (n indicates the number of eggs imaged per condition; 3 or more independent experiments conducted per condition; asterisks indicate a statistically significant difference compared with wildtype eggs; ****, P < 0.0001, two-tailed Fisher’s exact test). WT, wildtype; PB, polar body.

**Figure S4.**
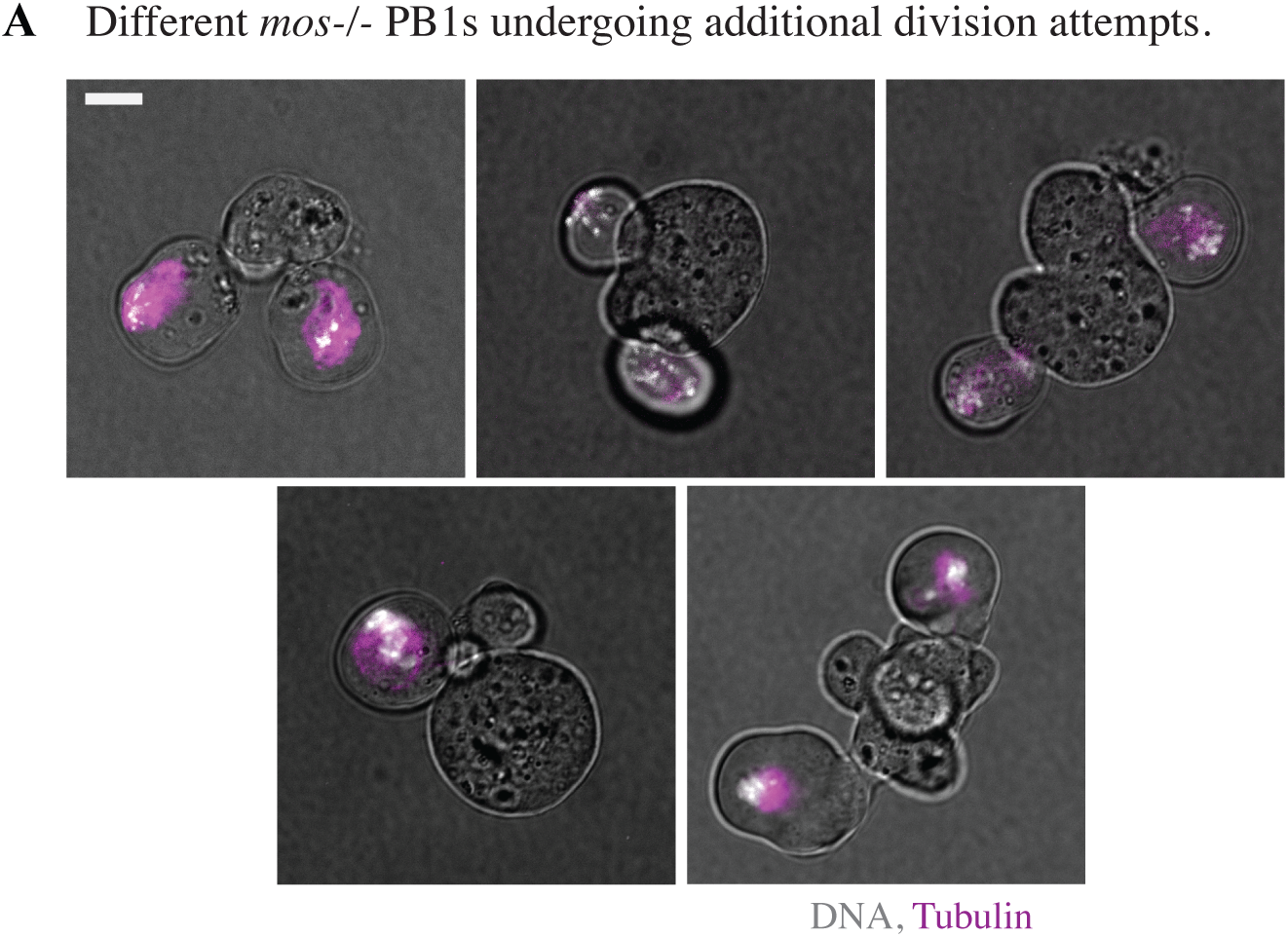
Isolated *mos^-/-^*polar bodies exhibit additional cell division attempts. **(A)** Individual images from time-lapse images of isolated *mos^-/-^* PB1 with abnormal morphological changes. Scale bars, 10µm. PB1, first extruded polar body.

## REFERENCES

1. Bellido-Quispe, D. K., Arcce, I. M. L., Pinzón-Osorio, C. A., Campos, V. F. and Remião, M. H. (2024). Chemical activation of mammalian oocytes and its application in camelid reproductive biotechnologies: A review. Anim Reprod Sci 266, 107499.

2. Cairo, G., Kholod, O., Palmer, O., Meytin, S., Goods, B. A. and Lacefield, S. (2025). Disrupted MOS signaling alters meiotic cell cycle regulation and the egg transcriptome. Reproduction 170, e250156.

3. Cao, L.-R., Jiang, J.-C. and Fan, H.-Y. (2020). Positive Feedback Stimulation of Ccnb1 and Mos mRNA Translation by MAPK Cascade During Mouse Oocyte Maturation. Front Cell Dev Biol 8, 609430.

4. Chaigne, A., Campillo, C., Gov, N. S., Voituriez, R., Azoury, J., Umaña-Diaz, C., Almonacid, M., Queguiner, I., Nassoy, P., Sykes, C., et al. (2013). A soft cortex is essential for asymmetric spindle positioning in mouse oocytes. Nat Cell Biol 15, 958–966.

5. Chaigne, A., Campillo, C., Gov, N. S., Voituriez, R., Sykes, C., Verlhac, M. H. and Terret, M. E. (2015). A narrow window of cortical tension guides asymmetric spindle positioning in the mouse oocyte. Nat Commun 6, 6027.

6. Choi, T., Fukasawa, K., Zhou, R., Tessarollo, L., Borror, K., Resau, J. and Vande Woude, G. F. (1996). The Mos/mitogen-activated protein kinase (MAPK) pathway regulates the size and degradation of the first polar body in maturing mouse oocytes. Proc Natl Acad Sci U S A 93, 7032–7035.

7. Colledge, W. H., Carlton, M. B., Udy, G. B. and Evans, M. J. (1994). Disruption of c-mos causes parthenogenetic development of unfertilized mouse eggs. Nature 370, 65–68.

8. Deng, M., Williams, C. J. and Schultz, R. M. (2005). Role of MAP kinase and myosin light chain kinase in chromosome-induced development of mouse egg polarity. Dev Biol 278, 358–366.

9. Dupré, A., Haccard, O. and Jessus, C. (2011). Mos in the oocyte: how to use MAPK independently of growth factors and transcription to control meiotic divisions. J Signal Transduct 2011, 350412.

10. Fan, W., Homma, M., Xu, R., Kunii, H., Bai, H., Kawahara, M., Kawaguchi, T. and Takahashi, M. (2021). The use of a two-step removal protocol and optimized culture conditions improve development and quality of zona free mouse embryos. Biochem Biophys Res Commun 577, 116– 123.

11. Hashimoto, N., Watanabe, N., Furuta, Y., Tamemoto, H., Sagata, N., Yokoyama, M., Okazaki, K., Nagayoshi, M., Takeda, N. and Ikawa, Y. (1994). Parthenogenetic activation of oocytes in c-mos-deficient mice. Nature 370, 68–71.

12. Hörmanseder, E., Tischer, T. and Mayer, T. U. (2013). Modulation of cell cycle control during oocyte-to-embryo transitions. EMBO J 32, 2191–2203.

13. Horner, V. L. and Wolfner, M. F. (2008). Transitioning from egg to embryo: triggers and mechanisms of egg activation. Dev Dyn 237, 527–544.

14. Hu, C.-K., Coughlin, M. and Mitchison, T. J. (2012). Midbody assembly and its regulation during cytokinesis. Mol Biol Cell 23, 1024–1034.

15. Jellerette, T., He, C. L., Wu, H., Parys, J. B. and Fissore, R. A. (2000). Down-regulation of the inositol 1,4,5-trisphosphate receptor in mouse eggs following fertilization or parthenogenetic activation. Dev Biol 223, 238–250.

16. Jiao, G., Lian, H., Xing, J., Chen, L., Du, Z. and Liu, X. (2022). MOS mutation causes female infertility with large polar body oocytes. Gynecol Endocrinol 38, 1158–1163.

17. Jung, G. I., Londoño-Vásquez, D., Park, S., Skop, A. R., Balboula, A. Z. and Schindler, K. (2023). An oocyte meiotic midbody cap is required for developmental competence in mice. Nat Commun 14, 7419.

18. Kalous, J., Tetkova, A., Kubelka, M. and Susor, A. (2018). Importance of ERK1/2 in Regulation of Protein Translation during Oocyte Meiosis. Int J Mol Sci 19, 698.

19. Kim, H. M., Kang, M. K., Seong, S. Y., Jo, J. H., Kim, M. J., Shin, E. K., Lee, C. G. and Han, S. J. (2023). Meiotic Cell Cycle Progression in Mouse Oocytes: Role of Cyclins. Int J Mol Sci 24, 13659.

20. Krauchunas, A. R. and Wolfner, M. F. (2013). Molecular changes during egg activation. Curr Top Dev Biol 102, 267–292.

21. Lefebvre, C., Terret, M. E., Djiane, A., Rassinier, P., Maro, B. and Verlhac, M.-H. (2002). Meiotic spindle stability depends on MAPK-interacting and spindle-stabilizing protein (MISS), a new MAPK substrate. J Cell Biol 157, 603–613.

22. Miao, Y., Ma, S., Liu, X., Miao, D., Chang, Z., Luo, M. and Tan, J. (2004). Fate of the first polar bodies in mouse oocytes. Mol Reprod Dev 69, 66–76.

23. Mihajlović, A. I. and FitzHarris, G. (2018). Segregating Chromosomes in the Mammalian Oocyte. Curr Biol 28, R895–R907.

24. Pines, J. (2011). Cubism and the cell cycle: the many faces of the APC/C. Nat Rev Mol Cell Biol 12, 427–438.

25. Salic, A. and Mitchison, T. J. (2008). A chemical method for fast and sensitive detection of DNA synthesis in vivo. Proceedings of the National Academy of Sciences 105, 2415–2420.

26. Sha, Q.-Q., Dai, X.-X., Dang, Y., Tang, F., Liu, J., Zhang, Y.-L. and Fan, H.-Y. (2017). A MAPK cascade couples maternal mRNA translation and degradation to meiotic cell cycle progression in mouse oocytes. Development 144, 452–463.

27. Terret, M. E., Lefebvre, C., Djiane, A., Rassinier, P., Moreau, J., Maro, B. and Verlhac, M.-H. (2003). DOC1R: a MAP kinase substrate that control microtubule organization of metaphase II mouse oocytes. Development 130, 5169–5177.

28. Uraji, J., Scheffler, K. and Schuh, M. (2018). Functions of actin in mouse oocytes at a glance. J Cell Sci 131, jcs218099.

29. Verlhac, M. H., Kubiak, J. Z., Weber, M., Géraud, G., Colledge, W. H., Evans, M. J. and Maro, B. (1996). Mos is required for MAP kinase activation and is involved in microtubule organization during meiotic maturation in the mouse. Development 122, 815–822.

30. Verlhac, M. H., Lefebvre, C., Guillaud, P., Rassinier, P. and Maro, B. (2000a). Asymmetric division in mouse oocytes: with or without Mos. Curr Biol 10, 1303–1306.

31. Verlhac, M. H., Lefebvre, C., Kubiak, J. Z., Umbhauer, M., Rassinier, P., Colledge, W. and Maro, B. (2000b). Mos activates MAP kinase in mouse oocytes through two opposite pathways. EMBO J 19, 6065–6074.

32. Wakayama, T. and Yanagimachi, R. (1998). The First Polar Body Can Be Used for the Production of Normal Offspring in Mice1. Biology of Reproduction 59, 100–104.

33. Zhang, Y.-L., Zheng, W., Ren, P., Hu, H., Tong, X., Zhang, S.-P., Li, X., Wang, H., Jiang, J.-C., Jin, J., et al. (2021). Biallelic mutations in MOS cause female infertility characterized by human early embryonic arrest and fragmentation. EMBO Mol Med 13, e14887.

34. Zhang, Y.-L., Zheng, W., Ren, P., Jin, J., Hu, Z., Liu, Q., Fan, H.-Y., Gong, F., Lu, G.-X., Lin, G., et al. (2022). Biallelic variants in MOS cause large polar body in oocyte and human female infertility. Hum Reprod 37, 1932–1944.

